# CellStudio: a Modular, Tunable and Accessible Platform for Analysis of Growth Factors Secretions in Cell Cultures

**DOI:** 10.1101/2024.10.03.614356

**Authors:** Enrique Azuaje-Hualde, Naiara Lartitegui-Meneses, Juncal Alonso-Cabrera, Asier Inchaurraga-Llamas, Yara Alvarez-Braña, Marian Martínez de Pancorbo, Fernando Benito-Lopez, Lourdes Basabe-Desmonts

## Abstract

Traditional cell culture methods face significant limitations in monitoring cell secretions with spatial and temporal precision. Advanced microsystems incorporating biosensors have been developed to address these challenges, but they tend to lack versatility, and their complexity, along with the requirement for specialized equipment, limits their broader adoption. CellStudio offers an innovative, user-friendly solution that exploits the Printing and Vacuum Lithography combined with bead-based assays to create modular and tunable cell patterns surrounded by biosensors. This platform allows for high-resolution, spatially resolved analysis of secreted proteins, such as VEGF and FGF-2, while being easily implementable in standard laboratory settings. CellStudio’s design is compatible with conventional laboratory equipment, facilitating its integration into existing workflows without the need for extensive training or specialized tools. Validation experiments using mesenchymal stem cells (MSCs) and HeLa cells demonstrated that CellStudio can detect small secretion levels from small cell clusters with high sensitivity as well as analyze diffusion profiles, remarking the possibilities for studying cell behavior. By offering a standardized, cost-effective approach to detailed cellular analysis, CellStudio significantly enhances the capabilities of traditional cell culture techniques, with broad applications across biological and biomedical research.

## 1. Introduction

Cell culture is the standard tool for biological, biomedical, and pharmaceutical research due to its versatility and broad application [1,2]. However, conventional cell cultures face significant limitations in accurately monitoring cellular processes within controlled microenvironments [3–5]. One major challenge is the measurement of secreted proteins, such as cytokines, enzymes, and growth factors, which are crucial for cell communication, immune responses, tissue repair, and overall cell development. Traditional assays for measuring cell secretions, such as enzyme-linked immunosorbent assay (ELISA), typically involve collecting and transporting cell culture supernatants to another location for external analysis. These methods often require large numbers of cells and provide bulk measurements, which hide the spatial and temporal details of secretion events. [6]

To address these challenges, researchers have explored integrating biosensors within cell cultures. Recent advancements in micro– and nano-fabrication have led to the development of microsystems that position biosensors close to cells, as previously reviewed by us [7]. For instance, Wang et al. described a system involving single-cell entrapment in PDMS wells, with antibody barcodes on the PDMS upper wall to capture cell secretion, which can then be detached and analyzed through fluorescence microscopy [8]. Cedillo-Alcantar et al. reported a microfluidic device designed for the entrapment of single cells with a single bead inside microchambers, where the beads incorporate antibodies to capture cell secretion, enabling analysis through single-bead immunoassays and fluorescence microscopy [9]. Additionally, Ansaryan et al. and Liu et al. developed a nanoplasmonic system for real-time cell secretion analysis, where single cells and tumoroids are captured on top of a plasmonic substrate comprised of a gold layer with well-ordered nanoholes, utilizing a specialized spectrometric microscope for the analysis [10–12]. However, while these systems offer advanced capabilities, they introduce increased complexity from both a fabrication standpoint and an end-user perspective. Their use requires specialized expertise and lack versatility in accommodating diverse biological scenarios. Additionally, they are limited in the degree of cellular microenvironmental control that can be achieved [7].

In response to these challenges, we have developed the CellStudio platform, a modular and tunable tool designed to simplify the integration of biosensing capabilities into cell patterns for spatially resolved monitoring of cell secretion. Our goal is to create an accessible and versatile platform that is easy to fabricate, user-friendly, and compatible with standard cell research protocols and microscopy. CellStudio has been developed combining established technologies like Printing and Vacuum Lithography (PnVlitho [13]) with bead-based assays[14,15]. It allows for high-resolution, localized analysis of secreted factors by positioning functionalized microbeads in close proximity to cell clusters, enabling detailed monitoring of cellular secretions. The platform supports hundreds of individual cell clusters surrounded by polymeric microbeads, each acting as an independent replicate boosting statistical reliability and data collection efficiency. Furthermore, CellStudio’s modular design allows researchers to easily adjust the number of cells or types of biosensors and create diverse experimental conditions within a single well plate.

One of CellStudio’s major strengths is its simplicity. The platform is designed to be fully compatible with standard laboratory equipment and protocols, combining a basic modified well plate bottom with innovative integrated patterns of beads and cell adhesion proteins. This design makes it accessible and practical for direct use with existing research infrastructure, such as optical and fluorescence microscopes. The ease of fabrication and seamless integration represents a significant advancement in cell research technology, offering a cost-effective and versatile solution for detailed cellular studies across a wide range of scientific and industrial applications.

## 2. Results and Discussion

### 2.1. Fabrication of CellStudio Using Printing and Vacuum Lithography

CellStudio substrates are fabricated using a technique known as Printing and Vacuum Lithography (PnVlitho). This technique was developed by our group to combine hierarchical metal nanoparticles structures with cell adhesion patterns [13]. Now, we extended the use of PnVlitho to create an array of identical cell adhesion areas surrounded by biosensors. PnVlitho process, summarized in **Figure 1A**, begins with microcontact printing, a method that involves the dry-transfer of proteins from a microstructured polydimethylsiloxane (PDMS) stamp onto a substrate, such as glass. In this study’s CellStudio configuration, PDMS stamps with uniformly spaced pillars were used to create an array of protein dots on the substrate. These protein dots serve as precise cell adhesion points, laying the groundwork for organized cell patterning within the substrate. In this configuration, 250 dots were printed per stamp, divided in two distinct arrays.

**Figure 1.**
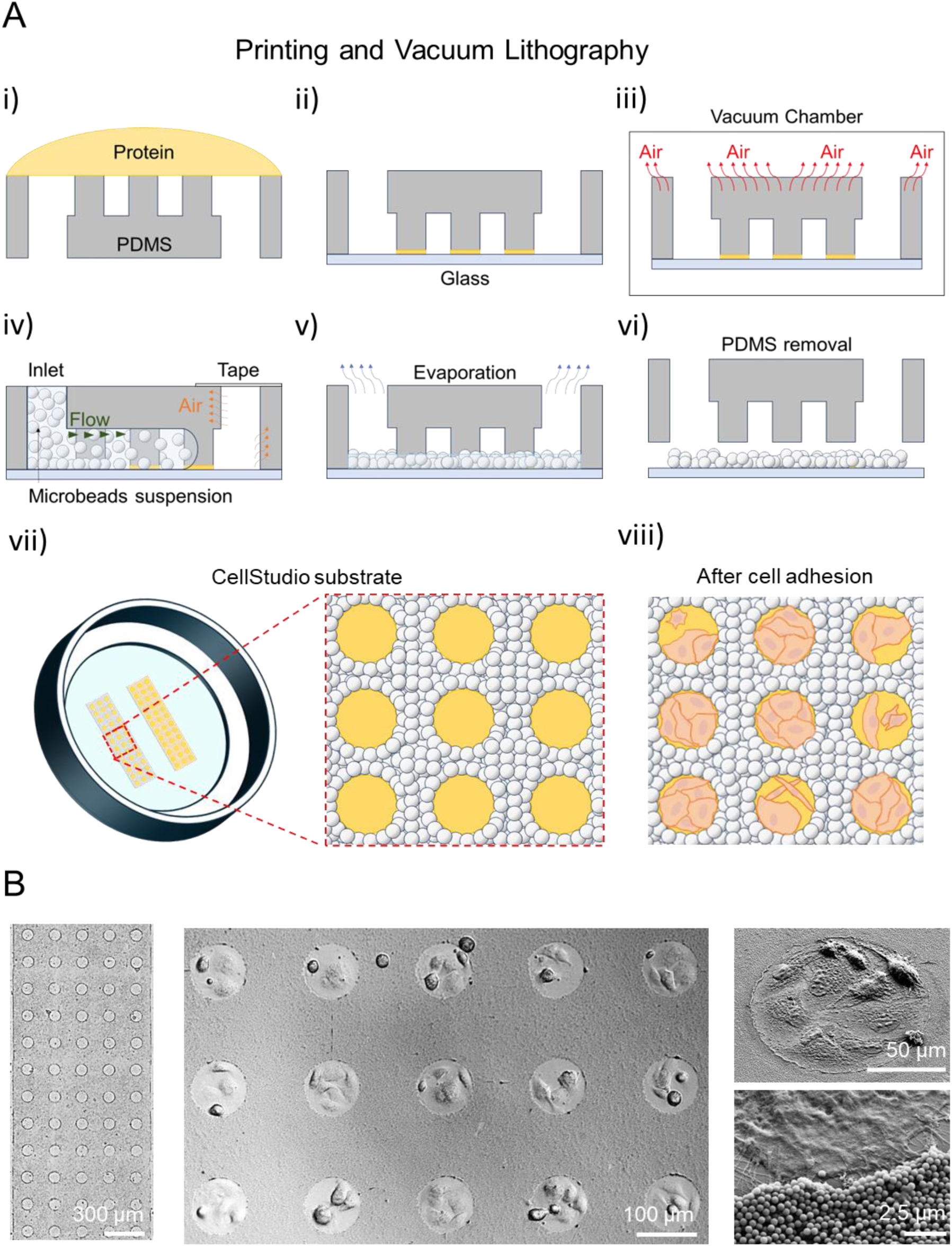
Fabrication of CellStudio substrates trough PnVlitho. A) Schematic representation of the Printing and Vacuum Lithography (PnVlitho) process: Polydimethylsiloxane (PDMS) stamps with pillars are incubated with a cell adhesion protein solution (i). After drying, the PDMS stamp is put into contact with the flat glass surface, transferring the protein to the substrate at the points of contact (ii). The PDMS stamps are then degassed in a vacuum chamber (iii). Upon returning to normal atmospheric pressure, the negative pressure generated by the absorption of air by the PDMS stamp allows the flow of a microbead suspension between its features (iv). After solvent evaporation (v), the PDMS stamp is detached (vi), providing the CellStudio substrate containing adhered microbeads surrounding the protein dots (vii). When incubated with cells, cells specifically adhere to each protein dot (viii). B) Brightfield (left and center) and SEM (right) images of cell loaded CellStudio substrates with increased zoom from left to right.

The next step in the PnVlitho process is vacuum-driven lithography. This stage takes advantage of the gas diffusion properties of PDMS to control the flow of a microbead suspension within the spaces between the PDMS stamp’s pillars. When the PDMS is placed under vacuum, the air within the polymer is removed. Once atmospheric pressure is restored, the PDMS draws air from all accessible spaces, including the areas between the carved features. Since the carved side of the PDMS stamp is sealed against the substrate, this negative pressure generates a controlled flow of a solution or suspension. By introducing a suspension of microbeads at this stage, the beads flow into the spaces around each PDMS pillar, forming a microbead pattern encircling the protein dots.

As the solvent evaporates and the PDMS stamp is removed, a two-dimensional pattern of cell adhesion proteins, encircled by a three-dimensional arrangement of microbeads, remains on the substrate. When this patterned substrate is exposed to a cell suspension, cells adhere specifically to the protein dots, forming individual cell clusters, each surrounded by microbeads. CellStudio substrates make cell loading and seeding simple by allowing to incubate cells directly on the platform without the need for additional steps, such as isolation of single cells using microwells or microfluidic traps [9,11,16,17].

Figure 1B shows a section of a CellStudio substrate, comprised of 60 dots homogeneously surrounded by streptavidin coated microbeads, after the incubation with a suspension of Mesenchymal Stem Cells (MSCs). Scanning Electron Microscopy (SEM) and optical bright field microscopy revealed a good homogeneity of the microbead pattern around the cell adhesion spots filled with cells. The images showed that cells on the substrate are in close proximity to the microbeads, what should be advantageous for accurate, sensitive, and efficient detection of secreted growth factors. The printed adhesive features are adaptable, allowing for precise regulation of cell capture, adhesion, distribution, confluence, and cell-cell contact by altering their shape, size or chemical composition [18–20]. The generation of hundreds of individual cell clusters on a single sample, each serving as an independent replicate, enhances the ability to obtain robust statistical data from the analysis.

The printed adhesive features are also customizable for the adhesion of various cell types. In addition to MSCs, three other distinct cell lines were tested: HeLa (human cervical cancer cells), MCF-7 (human breast cancer cells), and Jurkat (human T lymphocyte cells). Fibronectin was used to promote adhesion of adherent cells (MSCs, MCF-7, and HeLa), while a combination of fibronectin and anti-CD3 antibodies was employed to immobilize non-adherent cells (Jurkat). Patterning of all 4 cell types was successfully achieved, with both adherent and non-adherent cells forming distinct clusters on the biomolecule patterns, Figure 2A. Measurements from brightfield microscopy images revealed that the number of cells per protein dot was 4 ± 1, for MSCs, 12 ± 1 for HeLa, 6 ± 1 for MCF-7 and 14 ± 2 for Jurkat cells, Figure 2B.

**Figure 2.**
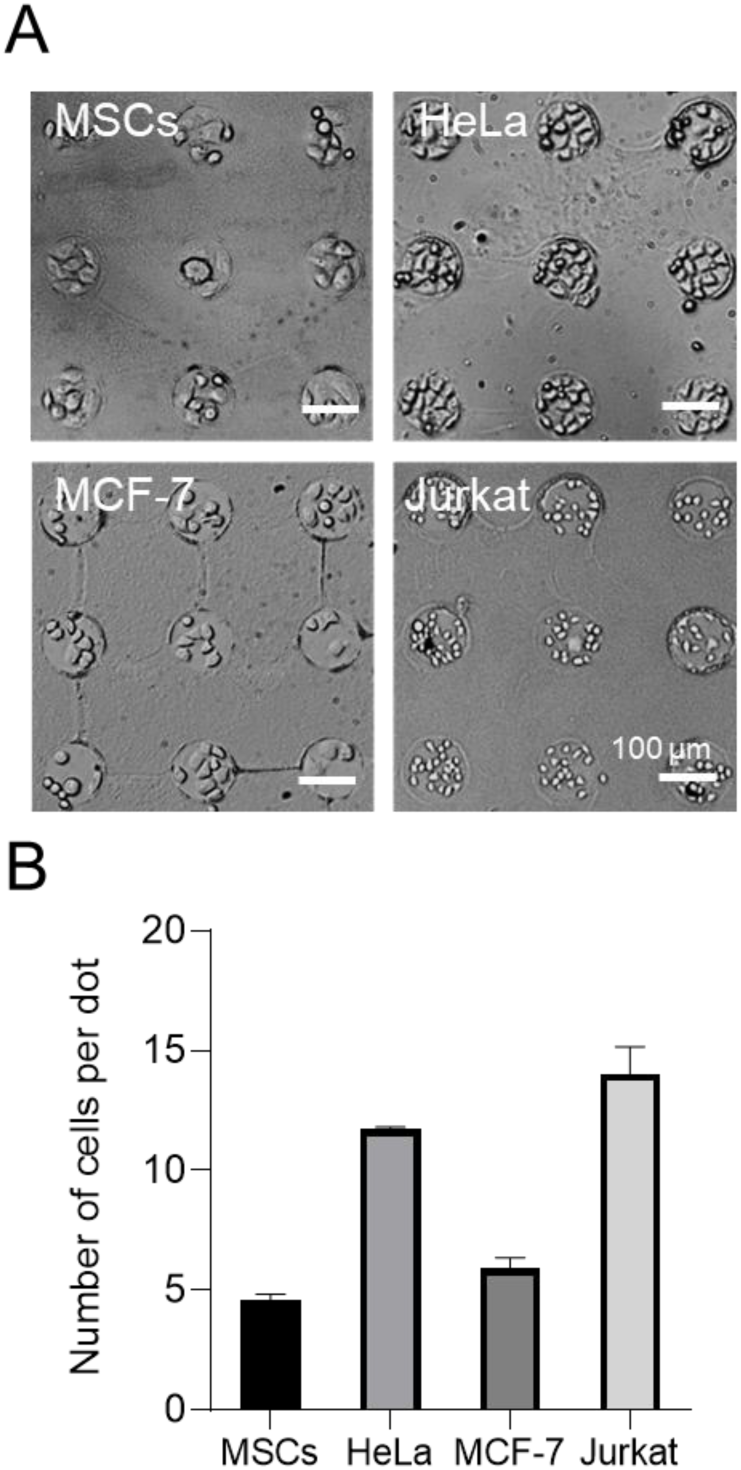
CellStudio loaded with different types of cells. A) Brightfield microscopy images of the patterned substrates containing different cell types, MSCs, HeLa, MCF-7 and Jurkat cells. B) Graphical representation of the number of cells per dot in each case. Error bars mean ± SED (n = 100 from 3 CellStudio patterns).

These results are consistent with the known properties of these cell types, demonstrating that the number of cells per dot is directly influenced by cell size and their tendency to spread. For instance, MSCs, which are relatively large (15-30 µm) and exhibit significant spreading on flat surfaces, had the lowest number of cells per dot. In contrast, Jurkat cells, which are smaller (10-15 µm) and do not spread, had the highest number of cells per dot. The cancer cell lines, HeLa and MCF-7, have sizes similar to MSCs but with less spreading capability, resulting in intermediate numbers of cells per dot. These results highlight the CellStudio platform’s effectiveness in accommodating a diverse range of cell lines.

### 2.2. Analysis of cell-secreted VEGF using CellStudio

Vascular Endothelial Growth Factor (VEGF), a growth factor known to induce angiogenesis and vasculogenesis, plays a pivotal role in numerous physiological and pathological processes, including wound healing, cancer progression, and tissue remodeling [21–23]. The secretion of VEGF from Mesenchymal Stem Cells (MSCs) is especially significant as it contributes to the pro-angiogenic environment necessary for effective tissue repair and regeneration [24–27]. By positioning microbeads functionalized with antibodies against VEGF in the surroundings of MSCs clusters (4-5 cells), VEGF secretion can be captured and then analyzed through fluorescence microscopy techniques. This way, after immunostaining, it is possible to quantify the localized secretion in the surroundings of hundreds of individual cell clusters in a single sample.

To demonstrate the usability of the CellStudio to analyze VEGF secretion from MSCs first, we first established a calibration curve. A CellStudio substrate without cells was incubated with solutions of known concentrations of VEGF (ranging from 1 to 1000 ng mL^-1^). Fluorescence microscopy was employed to directly image the samples after the immunostaining. For data analysis, the mean fluorescence intensity of annular regions 10 µm wide around the dots was assessed.

As expected, the fluorescence intensity measured in each sample increased with increasing concentrations of VEGF, and a clear correlation between the fluorescence intensity and the concentration of VEGF was observed, Figure 3. The obtained data were fitted into a four-parameter logistic (4PL) equation and the limit of detection was determined to be 0.15 ng mL^-1^. The obtained sensitivity level is comparable to the sensitivity of the gold standard conventional Enzyme-Linked Immunosorbent Assay (ELISA) [28]. The results indicate the potential of CellStudio as a tool for the detection of biomolecules upon successful capture of proteins in solution.

**Figure 3.**
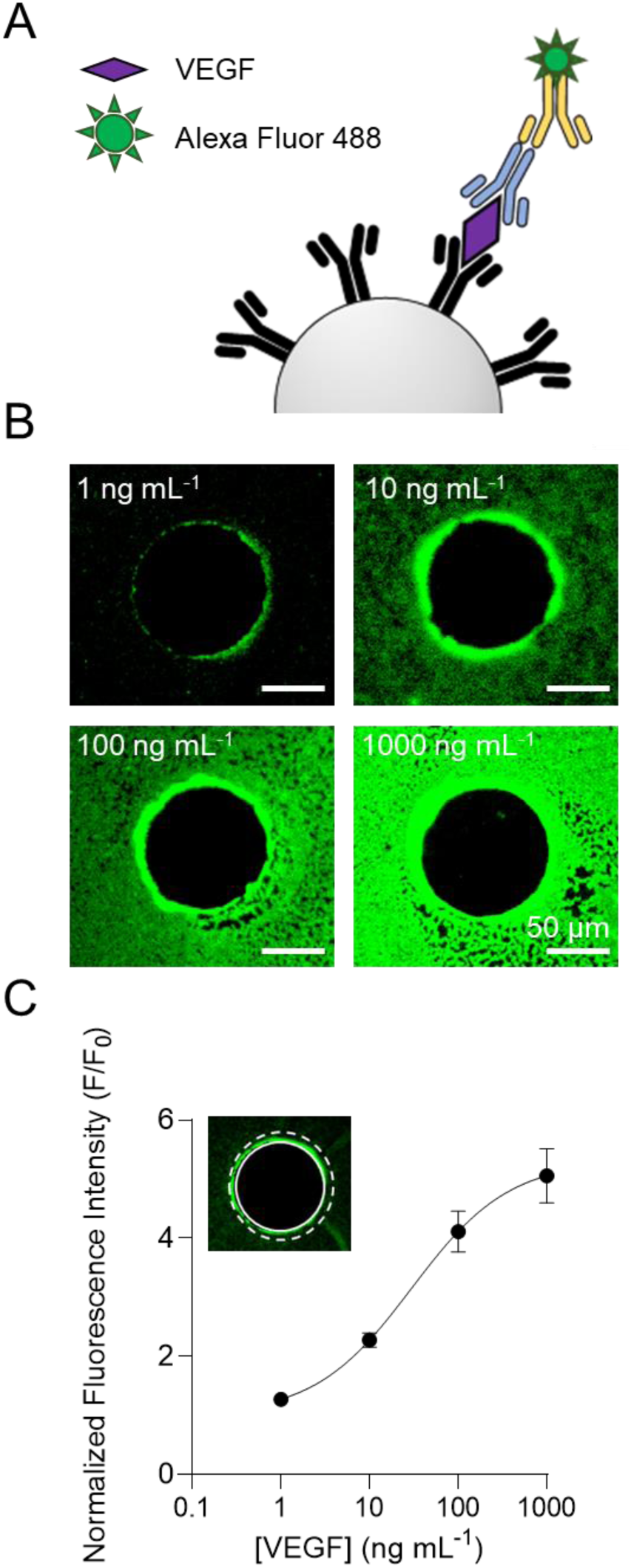
Calibration curve of VEGF detection on CellStudio substrates functionalized with VEGF immunoassay. A) Schematic illustration of the fluorescent immunoassay on a bead for VEGF detection. B) Fluorescence microscopy images of CellStudio substrates incubated with solutions of increasing concentrations of VEGF ranging from 1 to 1000 ng mL^-1^ and subsequent immunostaining. C) Plot of the fluorescence intensity of the images in a region of interest around each dot (10 µm annular region from the edge of the dots). Data is normalized to F_0_ corresponding to the mean value of the negative control (0 ng mL^-1^). Error bars represent mean values ± SD (n = 40 dots from 3 patterns).

After establishing a calibration curve, MSCs were patterned on a CellStudio substrate to create hundreds of distinct cell clusters, each consisting of 4 to 5 cells and positioned on 100 µm diameter fibronectin dots. Patterned cells were kept on the substrates over a 48-hour period allowing for VEGF secretion. VEGF was quantified using the previously calibrated fluorescent immunoassay.

Microscopy images clearly revealed positive immunostaining around the cell clusters, demonstrating that the cells secreted VEGF, which was effectively captured by the functionalized microbeads located nearby (Figure 4A). For data analysis, the mean fluorescence intensity of annular regions 10 µm wide around 65 different cell clusters was assessed. The mean fluorescence intensity around each cluster was 80 % higher than the negative controls (n = 65 clusters) Figure 4B. This intensity, based on the prior calibration, corresponded to a localized VEGF concentration of 5 ± 2 ng mL⁻¹ and a cell secretion rate of 1.7 to 2 ng per 10⁶ cells per day. These results were in agreement with the measurements obtained by an ELISA assay analyzing VEGF secretion from the supernatants of cultures of our MSC, as detailed in **Supporting Information 1**.

**Figure 4.**
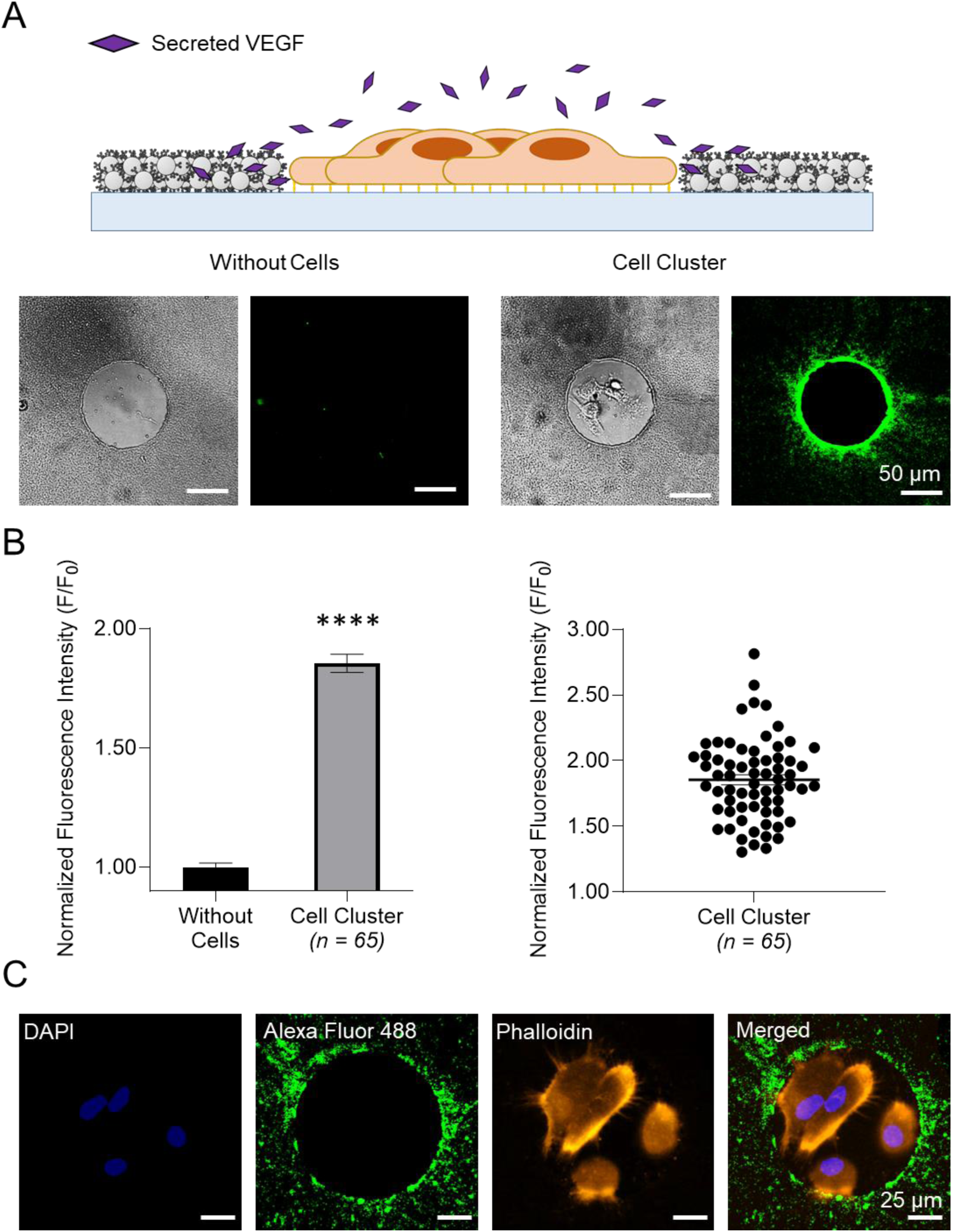
Detection of VEGF secreted from patterned MSCs clusters on CellStudio substrates functionalized with VEGF immunoassay. A) Top; Schematic cartoon representing a VEGF secreting cell cluster surrounded by the microbeads pattern functionalized with anti-VEGF antibodies. Bottom; Brightfield and fluorescence microscopy images of CellStudio substrates containing microbeads functionalized with anti-VEGF antibody, without cells (left) and with MSC clusters (right) after 48 hours of incubation. B) Bar plot and dispersion plot of the fluorescence intensity of annular regions of interest (ROIs) around each cell cluster (10 µm wide ring from the edge of cell clusters, n = 65 cell clusters from 3 patterns). Data was normalized to substrates F_0_ corresponding to the mean value of the negative controls (substrates Without Cells). Statistical significance was assessed using the non-parametric Mann-Whitney test (****p < 0.0001). C) Fluorescence microscopy images of a MSC clusters on CellStudio substrates containing microbeads functionalized with anti-VEGF antibody, showing cytoskeleton staining (Phalloidin, orange), nucleus staining (DAPI, blue) and secreted VEGF staining (Alexa Fluor 488 secondary antibody, green).

The ability to measure VEGF secretion from just 4 to 5 cells per dot highlights CellStudio’s high sensitivity and precision unlike conventional ELISA kits, which typically require a large number of cells to detect meaningful signal. The consistency across multiple clusters shows how effective CellStudio is in providing reliable data for studying cell secretion.

As in classical 2D cell culture studies, where researchers commonly assess cell morphology, proliferation, viability and intracellular structures through microscopy, CellStudio builds on these methods by enabling high-resolution observation of both cell morphology and cellular secretions. This was demonstrated by fluorescence images from CellStudio substrates, which reveal detailed visualization of cell nuclei using DAPI, cytoskeleton organization with phalloidin, and VEGF secretion with Alexa-488 labeled antibodies, Figure 4C. This combined approach simplifies the experimental process, cutting down on the need for multiple separate assays and making data collection and analysis more efficient.

Next, the CellStudiós capabilities to study the diffusion of secreted soluble factors was evaluated. For VEGF diffusion analysis, the gradient of fluorescence intensity was measured from the edge of the cell cluster up to 25 µm away. Fluorescence analysis revealed that secretion was specially concentrated majorly within the immediate 5 µm around each cell cluster, with a gradual decrease in intensity as the distance from the cluster increased, Figure 5. The image analysis showed that most VEGF secretion is localized near each cell cluster, with the observed fluorescence primarily reflecting secretion from the corresponding cluster. Beyond 10 µm, fluorescence intensity stabilized, slightly exceeding negative controls but remaining below the statistical limit of detection (LOD), suggesting that while VEGF diffusion may diffuse over larger distances, it does at minimal concentrations. These results demonstrated that the CellStudio platform effectively captures the diffusion profile of secreted factors, such as VEGF from cell clusters, which could be used to study cell-cell communication by looking how protein diffusion affects cell behavior in nearby clusters.

**Figure 5.**
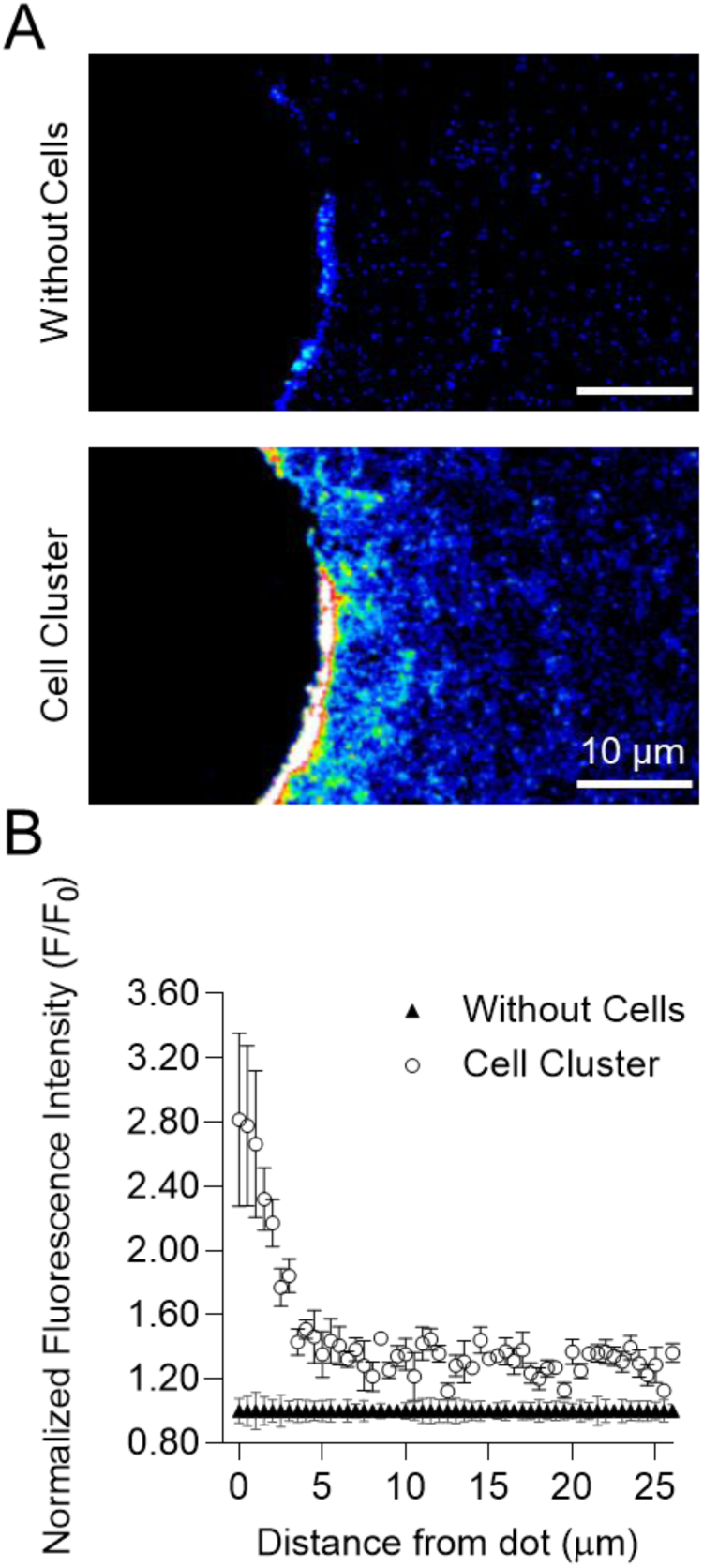
Analysis of secreted VEGF diffusion from MSC Clusters using the CellStudio platform. A) Fluorescence images (heat gradient scale) of dots in CellStudio substrates, without cells (top) and with cells (bottom), illustrating VEGF diffusion from the dot outward on CellStudio substrates functionalized with anti-VEGF antibody and stained with Alexa Fluor 488 secondary antibody after 48 hours incubation to allow secretion. B) Plot of the normalized fluorescence intensity around cell clusters, measured every 0.5 µm from the edge of the dot up to 25 µm. All data was normalized to F_0_ corresponding to the mean fluorescence value of the corresponding negative controls (substrates Without Cells). Error bars represent SEM (n = 10 dots from 3 different samples).

### 2.3. Evaluating CellStudio Platform for Comparative Analysis of Secretion of VEGF and FGF-2 by MSCs and HeLa Cells

Lastly, an experiment was designed to evaluate the sensitivity and specificity of the CellStudio platform in detecting and quantifying distinct growth factors, as well as to determine if the platform could provide insights into how different cell types influence their microenvironments through their secretions. The secretion of and FGF-2 by MSCs and HeLa cells were simultaneously analyzed and compared on identical CellStudio substrates. VEGF is essential for angiogenesis, promoting the formation of new blood vessels, while FGF-2 is critical for cell proliferation, differentiation, and tissue repair [29,30]. MSCs, known for their regenerative properties, secrete these growth factors to support tissue regeneration and angiogenesis. In contrast, HeLa cells, a well-established cancer cell line, secrete these factors to enhance their own growth and survival, reflecting their aggressive and proliferative nature [31]. Comparing these secretions provides insights into how regenerative versus cancerous cells influence their environments, which can deepen our understanding of cellular behavior in both health and disease.

For this experiment, we employed two distinct types of microbead patterns for detection: one using microbeads functionalized with biotinylated anti-VEGF antibodies, and another with microbeads functionalized with biotinylated anti-FGF-2 antibodies. The patterns of cell clusters surrounded by microbeads were incubated for 48 hours, followed by immunostaining and fluorescence microscopy imaging to evaluate the secretion levels. Fluorescence intensity measurements showed positive signals in the vicinity MSCs and HeLa clusters in both VEGF and FGF-2 biosensing patterns. The data showed notably higher secretion of VEGF and FGF-2 from the MSCs clusters compared to HeLa cells. The fluorescence intensity for the VEGF and FGF-2 detection substrates was 390 ± 20% and a 55 ± 7% higher for MSCs than for HeLa cells Figure 6.

**Figure 6.**
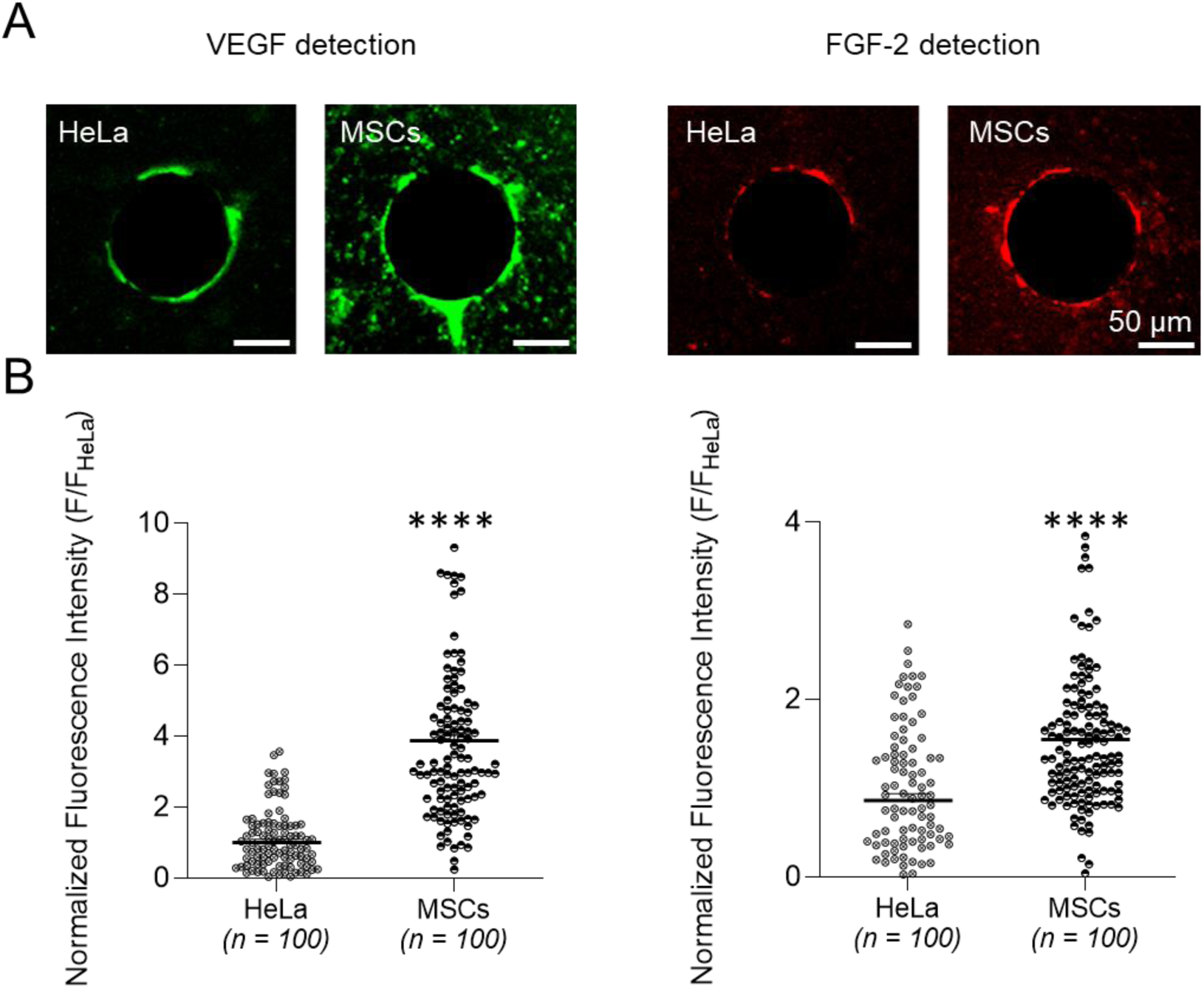
Comparative study of VEGF and FGF-2 secretion from MSCs and HeLa clusters. A) Fluorescence microscopy images of CellStudio substrates containing HeLa and MSCs clusters, after 48 h of incubation, to allow VEGF and FGF-2 secretion. VEGF secretion is shown in green and FGF-2 secretion is shown in red. B) Plot of the fluorescence intensity of a 10 μm wide annular ROI around each cell cluster on a VEGF (left) or a FGF-2 (right) sensing patterns after 48h incubation (n = 100 random cell clusters from 3 arrays in each case). All data was normalized to F_HeLa_ corresponding to the mean value of fluorescence intensities for the HeLa clusters for each protein. Statistical significance; non-parametric Mann-Whitney test (****p < 0.0001).

These results align with the distinct biological roles and characteristics of these cell types. MSCs are renowned for their regenerative properties and involvement in tissue repair and angiogenesis. Consequently, they secrete larger quantities of growth factors such as VEGF and FGF-2 to support tissue regeneration. HeLa cells, being cancerous, while also secrete VEGF and other factors to promote tumor growth and angiogenesis, often rely on the recruitment of MSCs to enhance tumor angiogenesis due to the latter’s higher secretory properties [31], explaining their reduced secretory capabilities.

Overall, these results demonstrate CellStudio’s suitability for comparative analysis of secretion from diverse cells on identical substrates. This capability enables comprehensive comparative studies of different cell types under standardized conditions.

## 3. Conclusions

This manuscript describes the platform technology CellStudio for cell analysis. By merging known techniques such as PnVlitho and bead-based assays it is possible to position microbeads functionalized with specific bioreceptors near to cells to enable the capture and quantification of secretions from individual cell clusters with minimal cell quantities. We have shown successful quantification of the VEGF secretion from cell clusters containing 4 or 5 cells, compatibility with standard protocols to study cells such as cellular staining, spatially resolved analysis of secreted growth factor diffusion and a comparative analysis of the secretion of VEGF and FGF-2 by two different cell types MSCs and HeLa cells.

For the manufacturing of CellStudio platforms, researchers only need a PDMS stamp, basic reagents such as beads and proteins, and a standard vacuum chamber. This simplicity allows for the creation of CellStudio substrates in standard research labs, lowering the barrier to entry for high-resolution cell analysis.

A major advantage of the platform is its compatibility with standard laboratory equipment and protocols, such as optical and fluorescence microscopy and immunostaining, allowing the measurement of intracellular events, including cytoskeleton dynamics, and cell morphology along with the monitoring of cell secretion. This integrated approach eliminates the need to transfer samples between different systems to acquire complementary data on cell behavior, thereby improving cost efficiency, and data consistency.

Finally, CellStudio is both tunable and modular, it may be tailored for different types of assays and cell types by modifying the size of the protein patterns and the composition of surrounding microbeads. On top of that, it is possible to create hundreds of identical areas on a single substrate, and several different substrates in the same well plate. This supports simultaneous experiments and replicates, comparative studies and a wide range of experimental configurations enhancing data collection and high-throughput applications.

In conclusion, we believe that CellStudio is an innovative platform that enhances traditional cell cultures by enabling the study of cell secretions in a controlled, spatially resolved environment. Unlike more complex microsystems, CellStudio’s simplicity in design, fabrication, and use is a strategic advantage, making the platform accessible, cost-effective, and versatile. We believe that this simplicity supports broad adoption, great reliability, and easy integration into existing workflows while maintaining its focus on the core innovation of spatially resolved cell secretion analysis in controlled microenvironments.

## 4. Materials and Methods

### 4.1. Fabrication of CellStudio substrates: Co-patterning of microbeads and cell adhesion clusters by Printing and Vacuum lithography (PnVlitho)

Polydimethylsiloxane slabs (PDMS, Ellsworth adhesives, Spain) channel-like structures (1000 x 5000 x 13 µm, width x length x height) with pillars inside (100 µm diameter separated by 100 µm) were employed. In all cases, PDMS slabs channels were punched twice, in order to generate a 2 mm diameter inlet and a 1 mm outlet. The pillars inside of the channels were wetted with a 50 µg mL^-1^ fibronectin solution (Fisher Scientific, Spain) in Phosphate Saline Buffer for 30 min, for capture and adhesion of adherent cell types. Afterwards, the PDMS slabs were rinsed with distilled water, dried with compressed air and attached to the glass bottom plate of a custom made well plate made of polymethyl methacrylate (PMMA, Evonik Industries AG, Germany) (**Supporting Information 2**). Glass substrates were previously oxidized in an air plasma chamber (29.6 W during 60 s, PDC-002-CE BlackHole, France). The resulted assembly of PDMS slabs on glass were put under vacuum (0.7 mbar) inside of a desiccator for 20 min. Afterwards, the outlets were plugged with tape and 2 µL of functionalized microbeads suspension were loaded in the inlets. The outlet of each channel was required to avoid phase separation of the bead suspension during flow, which interfered with the distribution of the beads inside the channel. The closing of the outlet was required to maintain the passive pumping inside the channel. The suspension was let to flow until it started filling the outlet. Streptavidin coated polystyrene microbeads of diameters of either 200 and 500 nm were employed in all experiments at suspension concentrations of 6 10^11^ microbeads mL^-1^ and 6 10^10^ microbeads mL^-1^ respectively. For further characterization of microbeads patterns of different sizes, see **Supporting Information 3**.

The tape was removed from the outlet after 5 min, and the PDMS slabs-glass assembly were left overnight at 4 °C for solvent evaporation. Finally, the PDMS slabs were removed and the resulting microbeads patterns surrounding the fibronectin dots could be observed on the substrate by optical microscopy.

The PMMA wells, with the now patterned glass bottom, were loaded with 1 mL of blocking solution 5 % Bovine Serum Albumin (BSA, Sigma Aldrich, Spain), 10 % casein (Fisher Scientific, Spain), 0.2 % milk powder in Phosphate Saline Buffer (PBS, Sigma Aldrich, Spain) for 2 h as blocking agent to avoid non specific binding.

### 4.2. Functionalization of microbeads patterns

For VEGF quantification assays, 400 µL of a 5 µg mL^-1^ biotinylated polyclonal Goat anti-Vascular Endothelial Growth Factor IgG solution in PBS (R&D Systems, USA) was added to CellStudio substrates containing streptavidin coated microbeads patterns (500 nm diameter beads) generated as previously stated. The substrates were incubated with the solution for 1 h at room temperature protected from light to allow the antibody to bind to the microbeads pattern trough biotin-streptavidin binding. Solutions were removed from the wells, which were rinsed three times with PBS afterwards.

For the comparative analysis of secretion of VEGF and FGF-2 from MSCs and HeLa cells, microbead patterns were created using 200 nm diameter microbeads pre-functionalized with the desired antibodies. For microbeads functionalization, stock microbead suspensions (volumes between 10-25 µL) were centrifuged at 14000 rpm for 9 min on an Eppendorf centrifuge 5425 (Germany). Afterwards, the microbeads were resuspended with 60 µL of the corresponding functionalizing solution and kept on constant mixing for 1 h. Functionalizing solutions included: distilled water, for the blank beads, 5 µg mL^-1^ biotin Polyclonal Goat anti-Vascular Endothelial Growth Factor IgG solution (R&D Systems, USA), for the generation of anti-VEGF functionalized beads, and 5 µg mL^-1^ biotin Polyclonal Goat anti-Fibroblast Growth Factor IgG solution (R&D Systems, USA), for the generation of anti-FGF-2 functionalized beads. Then, the suspensions were centrifuged at 14000 rpm for 9 min. The functionalized microbeads were resuspended in distilled water in the necessary volume to obtain a desired final concentration of 6 10^11^ microbeads mL^-1^. For the comparative analysis of secretion of VEGF and FGF-2 from MSCs and HeLa cells CellStudio patterns were made with microbeads mixes, comprised of a 1:1 mixture of blank microbeads and either anti-VEGF or anti-FGF-2 beads.

### 4.3. Cell seeding and cell culture on CellStudio substrates

For mesenchymal stem cells patterning, hair Human Follicle Mesenchymal Stem Cells (MSCs, isolated from human donors’ follicles) were detached from the flasks and were resuspended in serum free Dulbecco’s Modified Eagle’s medium (SFM, Fisher Scientific, Spain) at a concentration of 10^5^ cells mL^-1^. The same protocol was adapted and carried out for each cell line tested (MCF-7, HeLa and Jurkat cells).

Then, the blocking solution was removed from the PMMA wells containing the CellStudio substrate and the wells were rinsed three times with PBS. 750 µL of the cell suspension were loaded into them. To ensure specific cell adhesion to the fibronectin dots, the wells were left inside the incubator (37 °C, 5 % CO_2_) on constant oscillation using a rocker (Vari-Mix steep angle rocker, Thermo Fisher) for a maximum time of 120 min. Afterwards, the medium was retrieved, and the wells were rinsed three times with PBS to wash out any non-attached cell. For cell maintenance during the course of the experiments, cell-loaded CellStudio substrates were incubated with 750 µL of SFM.

### 4.4. Cells secretion and immunostaining

CellStudio substrates in SFM were kept during 48h in an incubator at 37 °C in a 5% CO_2_ atmosphere to allow cells to secrete. Before immunostaining, after the secretion period, cells were fixed by incubation with a 4% paraformaldehyde (Panreac Quimica, Spain) for 10 min at room temperature.

Subsequently, for VEGF detection, the substrates were incubated with 400 µL of a 1 µg mL^-1^ Mouse Monoclonal anti-VEGF IgG (R&D Systems, USA) with 10 % Goat Serum (v/v), 0.2 % milk powder (w/v) in PBS for 45 min. Then, the samples were rinsed with PBS three times and incubated with a secondary antibody solution containing 1 µg mL^-1^ Goat anti-Mouse IgG Alexa Fluor 488 (Fisher Scientific, Spain) for another 20 min. Finally, the samples were rinsed three times with PBS and analyzed.

For FGF-2 detection, fixed samples were incubated with 400 µL of a 1 µg mL^-1^ Alexa Fluor 647 Rabbit Monoclonal anti-FGF-2 IgG (abcam, USA) solution in PBS with 10 % Goat Serum (v/v) and 0.2 % milk powder (w/v) for 45 min. Then, the samples were rinsed three times with PBS and analyzed.

### 4.5. Image and data analysis

For the characterization of microbeads patterns, CellStudio substrates were imaged through Scanning Electron Microscopy (SEM) using a HitachiS-4800 SEM microscope.

The number of cells in each cell cluster were manually quantified from brightfield microscopy images. For the analysis of the fluorescence intensity in the vicinity of each spot, an annular region of interest (ROI) was used. This ROI had an inner diameter of 100 µm and an outer diameter 110 µm, covering an area 10 µm wide from the edge of the protein spot. Brightfield and fluorescence microscope images were taken with a modified Nikon Eclipse TE2000-S inverted microscope (USA), with and adapted Andor Zyla sCMOS black and white camera (Oxford Instruments, UK) and lumencor laser for excitation and Quad EM filter: 446/523/600/677 with 4 TM bands: 446/34 + 523/42 + 600/36 + 677/28. Further fluorescence microscope images were taken with an Olympus IXplore SpinSR10 spinning disk adapted to a camara HAMAMATSU ORCA FusionBT and widefield filters Ex: BP 352-402 | Em: LP 409, Ex: BP 460-480 | Em: BP 495-540, Ex: BP 530-550 | Em: LP 575 and Ex: BP 565-585 | Em: BP 600-690.

Microscopy images were processed by FiJi/ImageJ software. Data and Statistical analysis were performed in Excel 2016 and GraphPad Prism 8.

## 5. Acknowledgements

We acknowledge funding support from “Ministerio de Ciencia y Educación de España” under grant PID2020-120313GB-I00 / AIE / 10.13039/501100011033 and funding support from Basque Government, under Grupos Consolidados with Grant No. IT1633-22, and “Departamento de Salud del Gobierno Vasco” Grant code CART-gel. EAH acknowledges funding from the Basque Government, Department of Education, for predoctoral fellowship 2016 and funding support from the University of the Basque Country and the Spanish Government under the program “Margarita Salas” funded by “European Union – Next Generation EU”. JAC acknowledges funding from program “Investigo” funded by European Union – Next Generation EU. The authors also thank for technical support provided by “Analytical and High-Resolution Microscopy in Biomedice” SGIker (UPV/EHU/ ERDF, EU). Authors wish to thank technical assistance from the ICTS “NANBIOSIS”, more specifically by the Drug Formulation Unit (U10) of the CIBER in Bioengineering, Biomaterials & Nanomedicine (CIBER-BBN). Authors thank Dr. Alberto Gorrochategui from “Clínica Dermatológica Ercilla (Bilbao)” for providing the hair follicles from where hHF-MSCs were extracted.

## 7. Table of Contents

CellStudio is an innovative platform for cell biology research enabling spatially resolved analysis of cell secretions. Combining Printing and Vacuum Lithography with bead-based assays, it offers an accessible, modular, user-friendly tool compatible with standard lab equipment. CellStudio allows high-sensitivity detection of secreted proteins like VEGF, facilitating detailed studies of cells in various biological contexts.

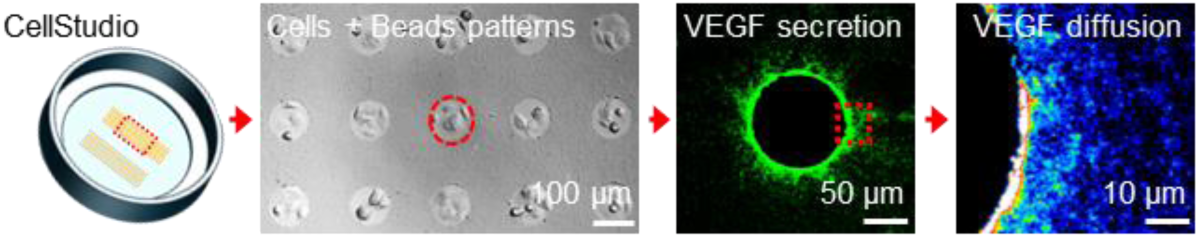

