## Supporting Information for "CellStudio: a Modular, Tunable and Accessible Platform for Analysis of Growth Factors Secretions in Cell Cultures"

**SI-1. Quantification of secreted VEGF from MSC cultures using ELISA kit**

The VEGF concentration in the supernatant of a conventional MSCs culture was quantified through an ELISA assay. MSCs were cultured in serum-free medium (SFM) in t75 flaks and were maintained in culture until reaching 80 % confluence. Afterwards, cells were detached and resuspended to a concentration of $10^5$ cells $mL^{-1}$. Suspension of cells were loaded in a 24 well-plate (500 µL per well) and were left in the incubator for 48 h to allow secretion of VEGF. Supernatants were retrieved, and cells were counted. Supernatants were then analyzed through conventional ELISA assays using a commercial kit (Quimigen, Spain). VEGF secretion was calculated for the ELISA assays and compared to the VEGF secretion values obtained by direct detection of MSCs patterns in CellStudio substrates **Figure SI-1 A**.

For quantification of cell secretion in CellStudio substrates, mean localized concentration of VEGF in the surroundings of each cell cluster was calculated using the calibration curve. For comparison between the different techniques, secretion was transformed as ng of VEGF secreted by $10^6$ cells in a single day using Equation 1, where *[VEGF]* refers to VEGF concentration (mg $mL^{-1}$) calculated, *V* refers to the volume (mL)**,** *nº of cells* refers to the mean number of cells found at the end of the assay and *day* the number of days cells were left secreting.

Eq. 1

$$\text{VEGF secretion} = \frac{[VEGF] \times V}{nº\ of\ cells\ \times days} \times 1000000$$

For the CellStudio substrates, *V* and *nº of cells* were calculated for individual cell clusters. In order to determine the *V* corresponding to the maximum volume in which VEGF may diffuse exclusively from each individual cell cluster, we defined a semi-sphere with a radius of 100 µm from the center of each dot, **Figure SI-1 B.** To calculate *nº of cells*, the mean number of cells per dot at the end of the assay was calculated.

The ELISA assay showcased a VEGF concentration 1.2 ± 0.2 ng $mL^{-1}$ of secreted VEGF after culturing the MSCs for 48 h (**Figure SI-1 C**). This corresponded to a cell secretion in the range of 2 – 2.5 ng per $10^6$ cells per day. This result present no significant differences with the values obtained from the direct analysis of VEGF secretion on patterned MSCs (1.7 – 2.1 ng per $10^6$ cells), validating CellStudio analytical potential.

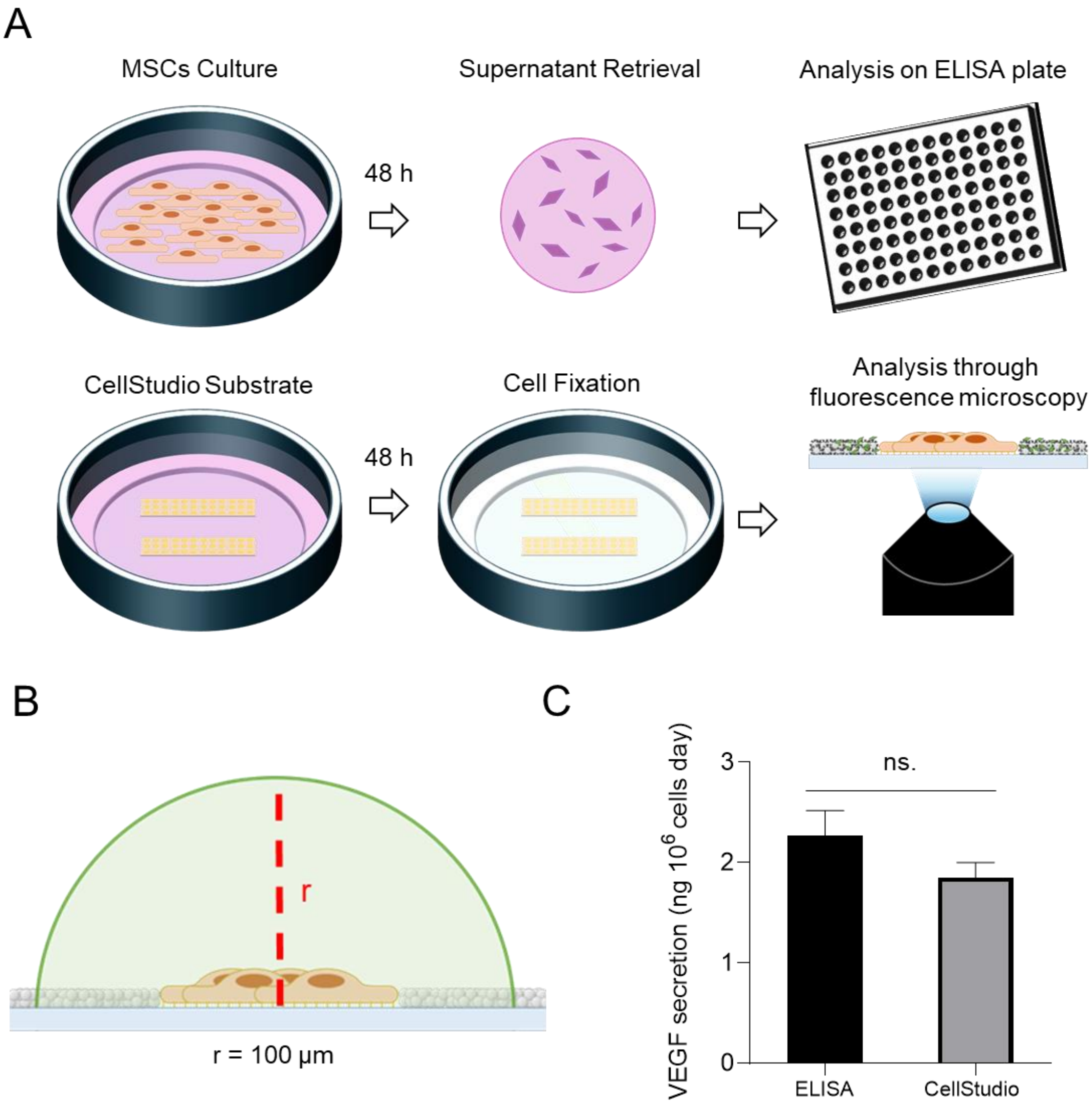


***Figure SI-1. Validation of VEGF detection.*** *A) Drawing of VEGF detection trough ELISA and CellStudio substrates. MSCs were cultured for 48 h on cell culture wells or in CellStudio substrate. Afterwards, for ELISA analysis, supernatants were retrieved and analyzed following commercial protocol. For analysis of CellStudio substrates, cells were fixed and analyzed through fluorescence microscopy after immunostaining. B) Drawing of dome-like volume defined as the volume for VEGF diffusion corresponding exclusively to each MSC cluster. C) Graphical representation of VEGF secretion (ng per* $10^6$ *cells per*

*day) secreted in MSCs supernatant, measured in ELISA as well as secreted by patterned MSCs clusters in CellStudio substrates functionalized with the immunoassay. Statistical significance: Mann-Whitney Test (ns. $p > 0.05$).*

### SI-1. Customized PMMA well plates

Customized two-well plates were fabricated by assembly of two 4 mm polymethylmethacrylate (PMMA) layers using a double side pressure sensitive adhesive (PSA) layer. Two-centimeter diameter wells were previously cut off the PMMA and the PSA substrates with a $CO_2$ laser (VERSA VLS2.30 Desktop Universal Laser System, USA). A glass cover slide served as the bottom plate of the wells, which was assembled with the PMMA wells using again a PSA film. **Figure SI-2**.

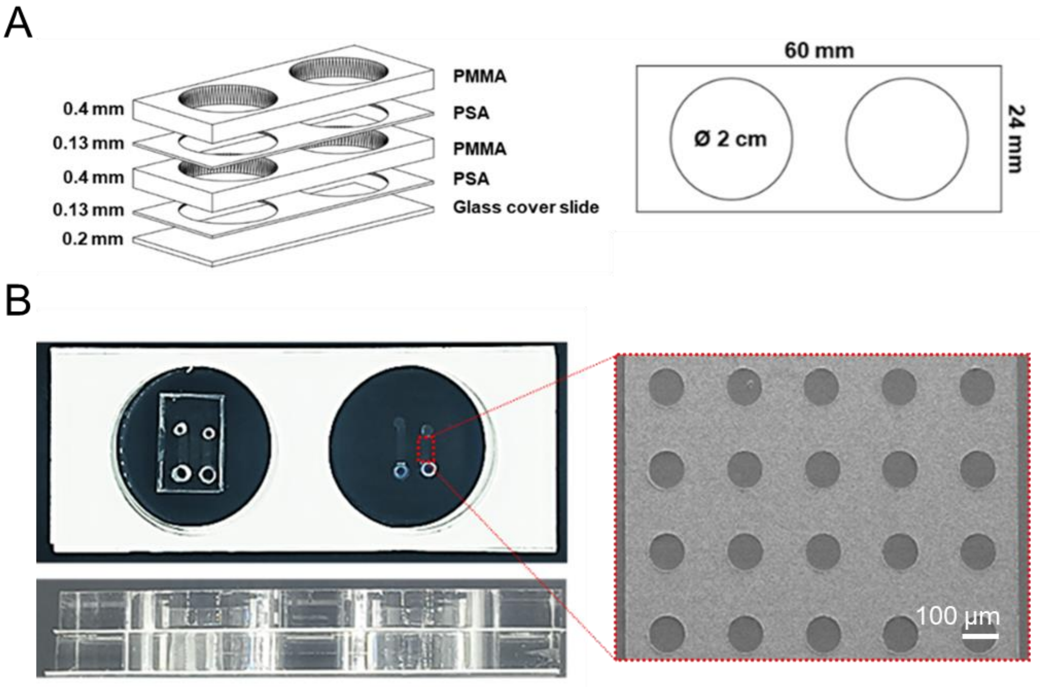


***Figure SI-2. CellStudio on custom made well-plate****. A) Schematic drawing of the PMMA culture well plate. B) Photographs of two well PMMA culture plate (top and side view), including two CellStudio substrates in each well, after (left well) and before (right well) the retrieval of the PDMS slab during the PnVlitho process. Zoom of the microbead pattern is a SEM image of microbeads surrounding the dots.*

**SI-3. Patterning of microbeads of different sizes and functionalizations.**

We evaluated the versatility of the CellStudio substrates making them with four different microbeads suspensions: 200 nm diameter streptavidin coated microbeads ($B_{beads}$), 500 nm diameter streptavidin coated $B_{beads}$, a 1:1 mixture of 200 nm diameter $B_{beads}$ and anti-VEGF funtionalized microbeads (VEGF-$ab_{beads}$) , and finally, a 1:1 mixture of 500 nm diameter $B_{beads}$ and VEGF-$ab_{beads}$. Characterization of the microbeads patterns by SEM showed that all four microbeads suspension produced a homogenous microbeads pattern throughout the surface and that 5 to 9 layers of microbeads were deposited over the glass surface in the areas between the adhesive dots (**Figure SI-3.1**)

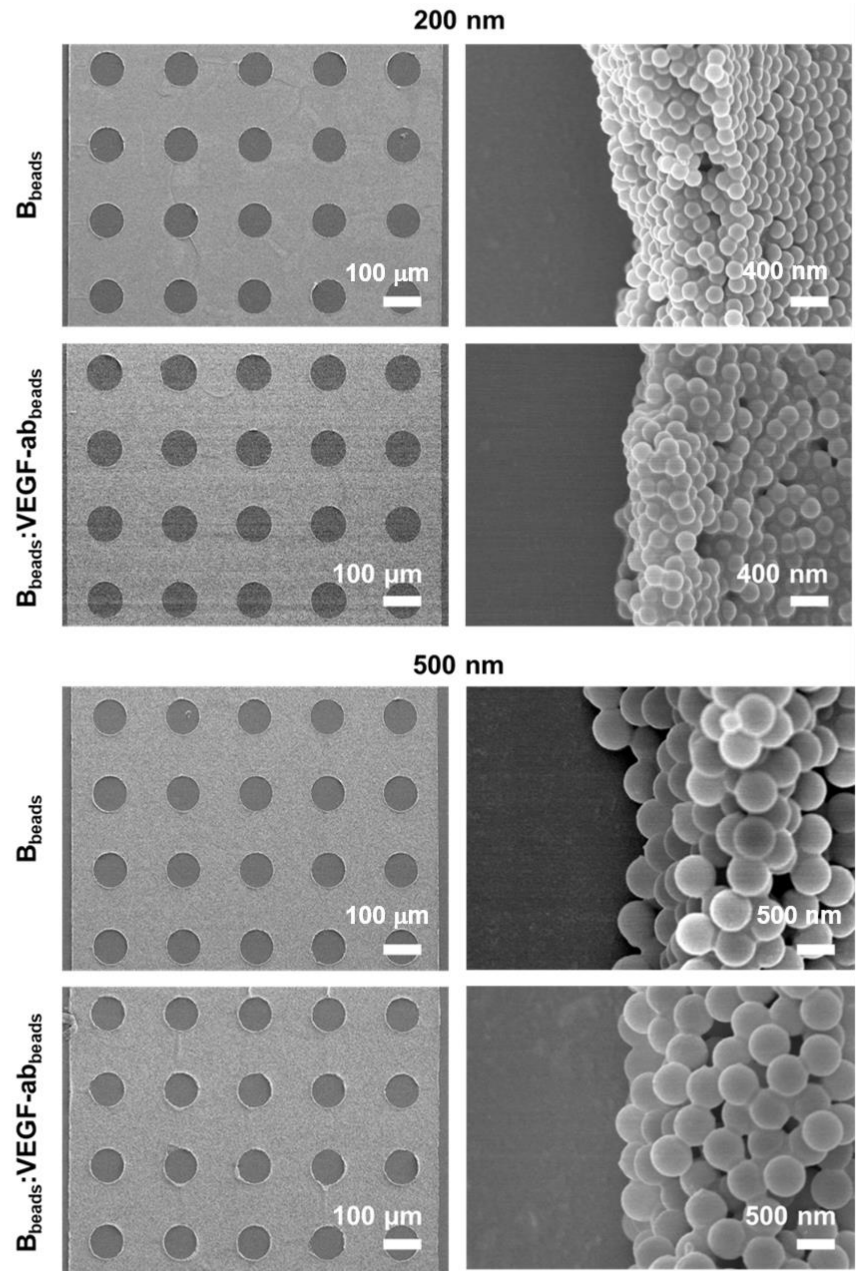


***Figure SI-3.1 CellStudio substrates varying size and functionalization of microbeads****. Scanning Electron Microscopy (SEM) images of microbeads patterns made with 200 and 500 nm diameter $B_{beads}$ (first row, third row) and mixes of $B_{beads}$:VEGF-$ab_{beads}$ of 200 and 500 nm diameter (second row, fourth row).*

Patterns of MSCs on CellStudio substrates comprised of fibronectin dots and 200 or 500 nm diameter $B_{beads}$ **Figure SI-3.2**

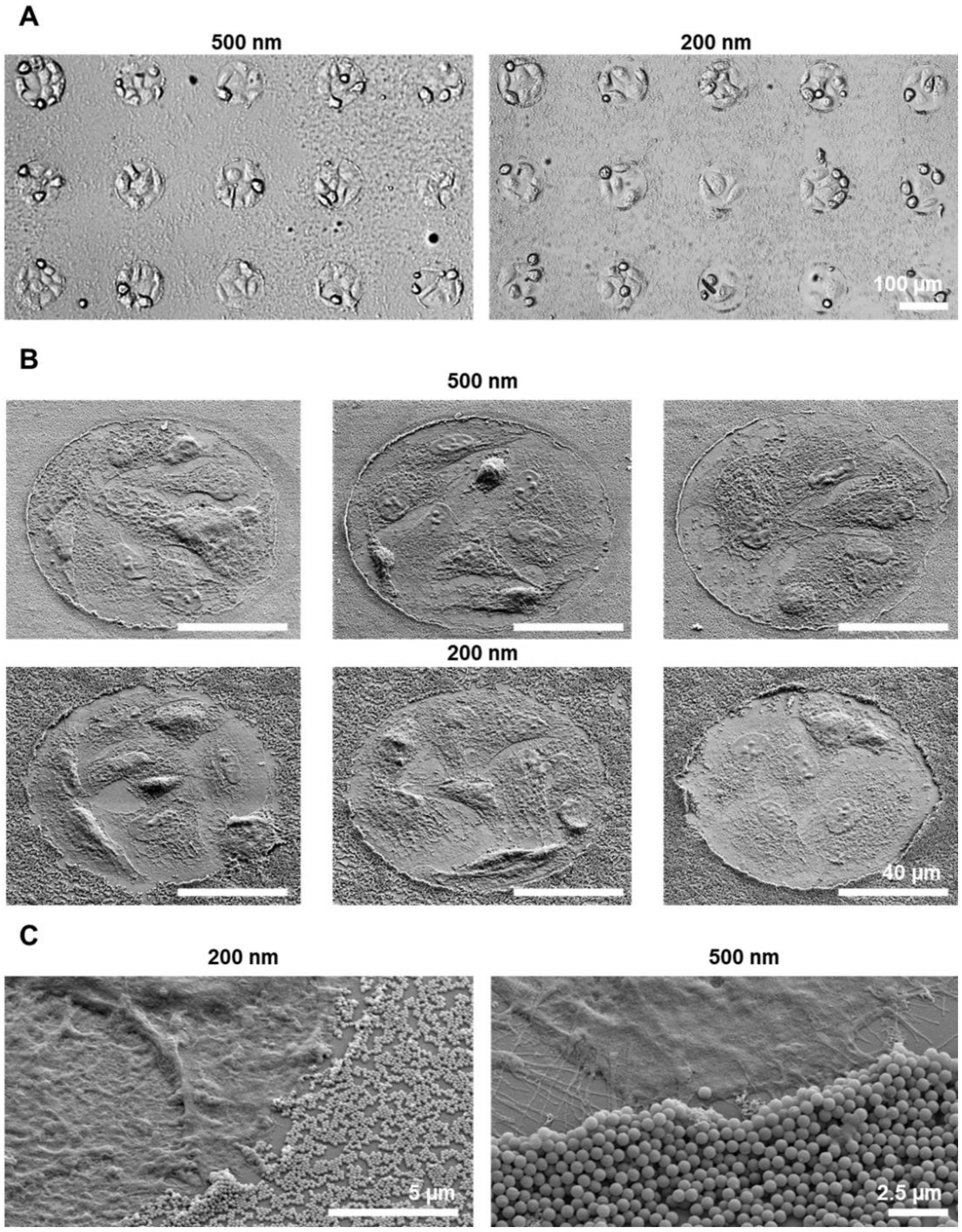


***Figure SI-2.2. MSCs loaded CellStudio substrates made with 200 and 500 nm diameters microbeads.*** *A) Brightfield microscopy images (A) and SEM images (B) MSCs loaded substrates made with 500 nm diameter (left) and 200 nm diameter (right)*

*$B_{beads}$ C) SEM images of higher magnification showing the interaction between the MSCs and the microbeads (200 nm diameter (left) and 500 nm diameter (right) microbeads).*
